# Percussive manipulations distinguish object play in capuchin and patas monkeys

**DOI:** 10.64898/2026.10.12.758487

**Authors:** Theresa Bier, Alice M.I. Auersperg, Remco Folkertsma, Isabelle B. Laumer

**Affiliations:** Department of General and Systematic Zoology, Institute of Biological Sciences, University of Rostock, 18059 Rostock, Germany; Comparative Cognition Unit, Messerli Institute, University of Veterinary Medicine Vienna, Vienna, Austria; Max-Planck-Institute of Animal Behavior, Development and Evolution of Cognition, Konstanz, Germany

**Keywords:** exploration, object play, tool-use, combinatory behavior, primate cognition

## Abstract

Object play has been proposed as a developmental precursor of flexible tool use. We applied a standardized object play protocol, previously used with corvids and parrots, to tufted capuchin monkeys (*Sapajus apella*), a member of a genus known for tool use, and patas monkeys (*Erythrocebus patas*), for which tool use has not been reported. Both groups were tested in a zoo across 15 sessions with a standardized set of objects and activity plates. All capuchins smashed objects, whereas only two patas monkeys did so, a difference consistent with the percussive foraging of capuchins. The species did not differ significantly in behavioral variability, object combinations, manipulation time, number of body parts used, or tail use, and combination rates were low in both. The species thus differed in a specific form of manipulation rather than in overall or combinatory object play. With one species per category and small samples, these results cannot establish whether combinatory object play is associated with tool use in primates.

## Introduction

Tool use provides a valuable window into the cognitive capacities, problem-solving strategies, and ecological pressures that shape behavior. Among our closest living relatives, habitual tool use in the wild is restricted to chimpanzees and orangutans, while only rarely observed in bonobos and gorillas (Breuer et al., 2005; Fox et al., 1999; Hare & Yamamoto, 2015; McGrew, 2010). Outside the great apes, habitual and spontaneous tool use has been documented in only very few primate taxa, including macaques (*Macaca* spp.) and baboons (*Papio* spp.) (Colbourne et al., 2021). Capuchin monkeys (*Sapajus* spp.) area striking exception among platyrrhines, which have otherwise been regarded as largely incapable or reluctant to use tools (Candland, 1987; Falótico, 2022; Westergaard & Suomi, 1993). In the wild and in captivity, capuchins use stones to pound and crack seeds and nuts, modify sticks to probe for food or water in cracks and insect nests, and use tool sets (e.g., Canale et al., 2009; Falótico & Ottoni, 2016; Ferreira et al., 2010; Fragaszy et al., 2005; Ottoni & Izar, 2008; Ottoni & Cardoso, 2014; Westergaard & Suomi, 1993).

How tool use develops during ontogeny remains a central question. Play has been proposed to provide opportunities to practice and refine motor and cognitive skills, thereby enhancing e.g. coordination, effectiveness, and problem-solving abilities (Bateson & Martin, 2013b; Nunes et al., 2004). Object play, in particular, may represent a foundational stage in the development of flexible tool use, as it may facilitate the acquisition of knowledge about object properties and refine the skills underlying problem-solving Graham & Burghardt (2010). Consistent with this view, infants and juveniles, and sometimes adults, of several species that habitually use tools and explore novel objects extensively through play (Jordan et al., 2021; Lambert et al., 2017; Ramsey & McGrew, 2005; Smith & Connolly, 1980; Tan, 2017). Object manipulation can be divided into primary actions, in which a single appendage handles an object, and secondary actions, which coordinate several appendages or bring an object into relation with another object or substrate (Hayashi et al., 2006; Hayashi & Matsuzawa, 2003; Takeshita, 2001; Torigoe, 1985). Secondary actions that combine an object with another object or with a substrate are considered key precursors of flexible tool use and technical problem-solving (Fragaszy & Adams-Curtis, 1992; Graham & Burghardt, 2010; Hayashi et al., 2006). In human infants, combinations of two free objects appear at the age of approximately eight months, combinations of objects with features of the environment, such as objects into cavities, at about ten months, and tool use directed at a goal after the second year (Connolly & Dalgleish, 1989; Greenfield et al., 1972; Rat-Fischer et al., 2012). There is some evidence that combinatory manipulation emerges before tool use during ontogeny in several species, including tufted capuchins, chimpanzees, orangutans, long-tailed macaques, baboons, and New Caledonian crows, although this evidence is correlational (reviewed in Colbourne et al. (2021))

Among non-human animals, object combinations have been studied predominantly in primates and some corvids and psittaciformes which exhibit object play behavior comparable to those of primates (Auersperg et al., 2011, 2014; Graham & Burghardt, 2010). Across primates, the extent of object manipulation behaviors increases from strepsirrhines through platyrrhines and catarrhine monkeys to great apes (Vauclair, 1984). In a survey of 74 primate species, Torigue (Torigoe, 1985) found complex motor actions and combinatory behaviors during object play only in capuchins and great apes, and he argued that the complexity of object combinations, rather than the overall diversity of object play, predicts the emergence of tool use. Great apes stand out for the diversity and complexity of their exploratory object manipulation, including bimanual manipulation, the coordinated use of multiple body parts as well as object-object and object-substrate interactions (Byrne et al., 2001; Heldstab et al., 2016; Laumer et al., 2025; Matsuzawa & Hayashi, 2003; Torigoe, 1985). For instance, Hayashi & Matsuzawa (2003) observed that inserting behavior emerges frequently and from an early age in infant chimpanzees tested on spontaneous object-object combinations. Captive capuchins (*Sapajus apella*) bang, throw, and rub objects and insert them into holes, wire mesh or containers (Visalberghi & Guidi (1998), and they combine objects with substrate during manipulation (Brunon et al. (2014). In the wild, capuchins of the genus Cebus also engage extensively in object play. White-faced capuchins (*Cebus capucinus*) interact with sand, water, and rocks, and explore the affordances of their environment independently from an early age (Perry et al. (2017). Object manipulation increases markedly in *Sapajus apella* and *Sapajus libidinosus* between one and six months of age (Araujo et al., 2024; Fragaszy & Adams-Curtis, 1997) and further between seven to twelve months (Fragaszy & Adams-Curtis, 1997). Comparable trends occur in humans and great apes, where early development is likewise marked by intensive exploratory object manipulation (Lamon et al., 2018; Laumer et al., 2025; Schuppli et al., 2021)

On an evolutionary scale, object play in a species’ repertoire may increase the likelihood that descendant species evolve tool use, regardless of the adaptive origin of play (Bateson & Martin (2013a)). Colbourne et al. (2021) proposed that an intrinsic motivation to combine objects underlies the onset of tool use directed at external objects, and that the form of a species’ combinations is shaped by its ecological niche. Species with high rates of functional object interactions or advanced motor abilities are therefore expected to play intensely with objects even in the absence of an immediate goal (Auersperg et al., 2014). Auersperg and colleagues tested this expectation with a standardized protocol in which subjects received wooden objects and activity plates with tubes, holes and poles. Across three corvid and three parrot species, the most complex combinations occurred in juvenile New Caledonian crows and in Goffin cockatoos. Both species have previously demonstrated sophisticated tool use: New Caledonian crows are the only corvids known to manufacture diverse foraging tools in the wild, while Goffin cockatoos show innovative stick tool use, hook bending, and tool set use (Auersperg et al., 2012; Hunt, 1996; Shumaker et al., 2011). By contrast, jackdaws, which do not rely on caching food or tool use, combined objects less often than both species. The type of combination also reflected ecology: insertions predominated in the crows, which extract food with tools, and play caching in ravens, which cache food. Age effects, however differed between species. Adult New Caledonian crows showed almost no interest in the objects, whereas combinatory behaviors of adult and juvenile Goffin cockatoos did not differ significantly (Auersperg et al., 2014). To our knowledge, this protocol has not been applied to primates, and it remains unknown whether primate species that differ in tool use show a comparable pattern.

We compared object play in captive tufted capuchin monkeys (*S. apella*) and patas monkeys (*Erythrocebus patas*), a cercopithecid species for which neither tool use nor pronounced object play has been reported. A zoo setting permits standardized observations, which is difficult to achieve in the wild. Following Auersperg et al. (2015) and Auersperg et al. (2014), we presented both species with a standardized set of objects scaled to body size and with activity plates, tested subjects in their social groups, and coded object play with a focus on combinations with other objects and substrates. Because the two species differ in phylogeny and ecology as well as in tool use, this design can identify species differences in object play but cannot attribute them to tool use alone. We tested the following predictions:

(I) As highly opportunistic omnivores capable of exploiting diverse habitats and environmental conditions (Fedigan, 2017), capuchin monkeys are predicted to exhibit greater variability in object play behaviors than the less opportunistic patas monkeys. In parrots, combinatory object play occurred in an Amazon species with a broad habitat range but not in a closely related species restricted to one habitat (Auersperg et al., 2015).
(II) Capuchins are predicted to display smashing behavior more frequently than patas monkeys. In birds, the type of object combination reflected foraging ecology (Auersperg et al., 2014), and capuchins rely on extractive foraging (de Andrade & de Sousa, 2018; Howard et al., 2018; Izawa & Mizuno, 1977).
(III) Capuchins are predicted to perform more complex object manipulations overall and interact more frequently with the activity plates than patas monkeys.
(IV) Juveniles of both species are expected to combine objects more often than adults, reflecting the decline of play with age (Pellis & Ham, 2024). Object combinations by adults are predicted to occur only in capuchins. Object play in adults has been proposed to support foraging in species that inhabit diverse or unpredictable environments (Diamond & Bond, 1999; Godfrey-Smith, 2001; Graham & Burghardt, 2010; Hall, 1998; Schuppli et al., 2017). In orangutans, adults from a population that use tools show m ore exploratory object combination than adults of a population that does not (Schuppli et al., 2017).
(V) Capuchins are assumed to spend more time manipulating objects than patas monkeys.
(VI) Both species are expected to use a similar number of body parts during object manipulation since they both regularly engage in terrestrial activity, allowing frequent use of all limbs for exploration (Laumer et al., 2025; Mittermeier et al., 2013; Visalberghi & Fragaszy, 2013). However, capuchins are predicted to show more tail use during object play than patas monkeys due to their prehensile tail, a functionally fifth extremity, which can be used during object manipulation than patas monkeys, whose tail is not prehensile (Ankel-Simons, 2007; Dunn & Cristóbal-Azkarate, 2016)

## Method

### Subjects and experimental history

The subjects of the study were one group each of patas and capuchin monkeys kept at Rostock Zoo. The patas group consisted of eight individuals (four adults, four juveniles) while the capuchin group consisted of six individuals (four adults, two juveniles; see Table 1). Both groups had access to two connected indoor enclosures (patas: 47 m^2^, capuchins: 60 m^2^) and a spacious outdoor enclosure (patas: 1270 m^2^, capuchins: 850 m^2^), each equipped with various climbing structures including tree trunks, wooden beams and ropes. Depending on weather conditions and outside temperature, animals could switch freely between indoor and outdoor enclosure or were kept indoor completely. All animals had ad libitum access to water and were fed several times a day with a variety of vegetables, fruits and seeds. The capuchin monkeys were housed together with two white-nosed coatis (*Nasua narica*), which were present in seven of the 15 test sessions, typically for less than 30 s and once for approximately 2min. They did not interact with the objects beyond sniffing them.

**Table 1:** Name, sex, age and age class of the participating patas (*Erythrocebus patas*) and capuchin monkeys (*Sapajus apella*) housed at Rostock Zoo.

| Individuum | Sex | Age (in years) | Age class | Species |
| --- | --- | --- | --- | --- |
| Asis | m | 1 | juvenile | *Erythrocebus patas* |
| Atu | m | 2 | juvenile |  |
| Nuru | m | 3 | juvenile |  |
| Njabi | m | 4 | juvenile |  |
| Zahina | f | 9 | adult |  |
| Jengo | m | 10 | adult |  |
| Zwena | f | 10 | adult |  |
| Uzima | f | 19 | adult |  |
| Tia | f | 3 | juvenile | *Sapajus apella* |
| Tica | f | 5 | juvenile |  |
| Mishki | f | 16 | adult |  |
| Avilex | m | 22 | adult |  |
| Santos | m | 23 | adult |  |
| Scharti | f | 38 | adult |  |

The patas monkeys had not participated in any behavioral test or cognition experiments before, whereas the capuchin monkeys participated in an experiment that investigated the ability to identify unknown food locations by excluding logical alternatives (inference by exclusion) using a standard two-cup test as part of the master’s thesis by Felix Albe (2024). None of the subjects had participated in experiments in the context of object play before and none of them has previously been presented with the objects used in this experiment.

### Ethics statement

Animal husbandry complied with the minimum requirements of the European Association of Zoos and Aquaria (EAZA) and the Mammal Expert Report for the accommodation and care of animals, and explicit ethical approval was obtained from Zoo Rostock. As the study was based strictly on non-invasive behavioral observations, it was not classified as an animal experiment under German law (§7 Bundestierschutzgesetz). Participation was voluntary, and animals could withdraw at any time; food and water were available throughout, and the animals’ health and welfare were monitored continuously before, during and after the experiments.

### Experimental setup

#### Apparatus

Similar as in the studies by Auersperg et al. (2014) and Auersperg et al. (2015), four different “activity plates” were prepared. These plates consisted of: (a) a 22 mm-thick plate with four holes drilled into its wooden base, (b) four dowel poles (c) four vertical tubes and (d) four horizontal tubes, each of varying diameters (see Figure 1). To improve stability and prevent the animals from moving the plates within their enclosures, two plates were secured to a larger wooden base, respectively.

**Figure 1:**
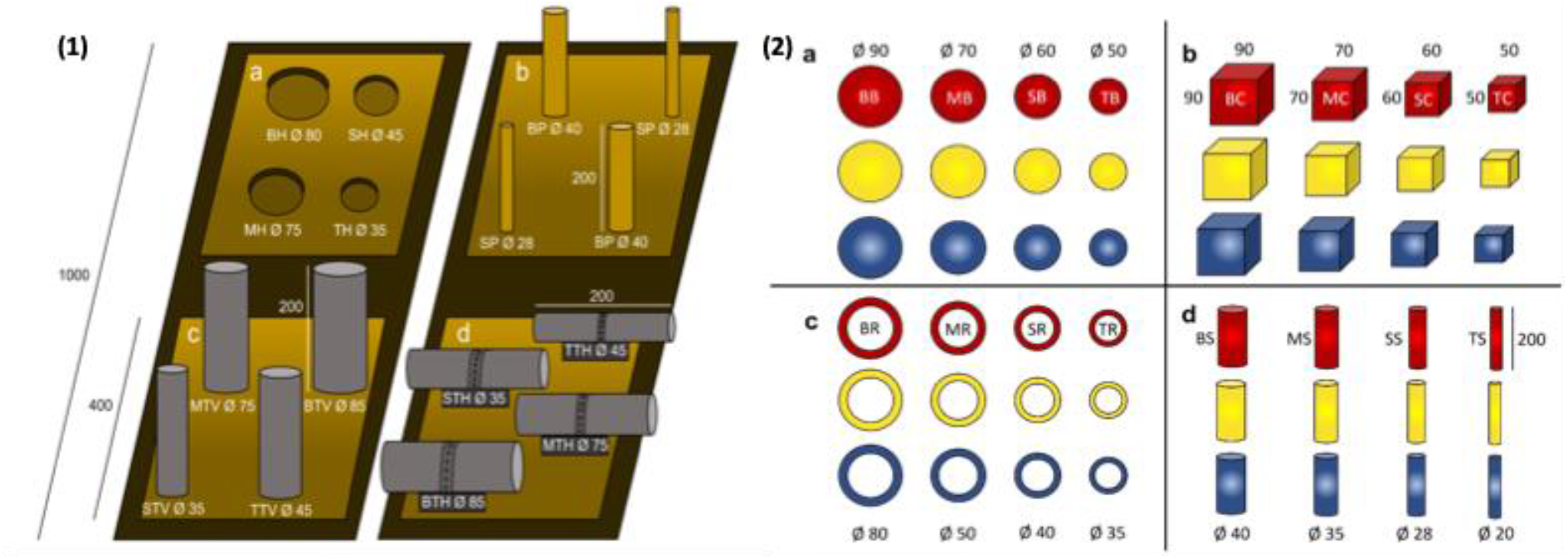
**(1)** Schematic layout of the activity plates (see SI, section A, Figure S1 for a photo). All dimensions in mm. **(a)** holes drilled into the wooden base (BH: big hole, MH: medium hole, SH: small hole, TH: tiny hole); **(b)** poles (BP: big pole, SP: small pole); **(c)** vertical tubes (BTV: big tube vertical, MTV: medium tube vertical, STV: small tube vertical, TTV: tiny tube vertical); **(d)** horizontal tubes (BTH: big tube horizontal, MTH: medium tube horizontal, STH: small tube horizontal, TTH: tiny tube horizontal). The length of poles and tubes was 200 mm, but due to the mounting design, the usable internal depth of the vertical tubes was only 150 mm. **(2)** Wooden objects used in the present study. All dimensions in mm. **(a)** balls (BB: big ball, MB: medium ball, SB: small ball, TB: tiny ball); **(b)** cubes (BC-TC); **(c)** rings (BR-TR); **(d)** sticks (BS-TS). The length of the sticks was 200 mm.

In addition, subjects were offered a set of wooden objects painted in three colors (red, blue and yellow) which should be clearly distinguishable by both patas and capuchin monkeys (Gomes et al., 2002; Jacobs et al., 1996; Macdonald et al., 2024; Pessoa et al., 1997). The objects, painted with child-safe, nontoxic acrylic paint, varied in four sizes and four shapes: balls, cubes, sticks and rings (see Figure 1).

#### Experimental procedure and video data acquisition

Video data was recorded at Rostock Zoo between 22 October and 18 December 2025. To avoid stress from social separation and to eliminate neophobia towards novel objects placed in the enclosure, which can affect play behavior (Torigoe, 1985), all subjects were tested in their usual housing and group settings and were presented with the activity plates prior to the first experimental session (habituation phase, see Appendix section A for details).

Immediately before testing, the activity plates were positioned in a rectangular configuration on the floor of the indoor enclosure, with approximately 30 cm of spacing between adjacent plates. The wooden objects were then randomly distributed in a semicircle around the plates, at a distance of about 30 – 40 cm depending on the enclosure size (also see Figure A1 in Appendix section A).

Each monkey group was tested in 15 sessions, each lasting 30 minutes, starting from the moment any individual first touched an object (also see Appendix section A for details). All object-directed behaviors were documented using two high-definition video cameras, positioned at two angles outside the enclosures at a distance of approximately 2 m. Additionally, object-directed interactions that could not be filmed were described verbally by the observer. If the animals did not touch any of the objects within 30 minutes, the session was terminated and repeated on another day; this occurred twice for the patas monkeys and once for the capuchins. Due to an injury, one patas monkey (Zahina) did not participate in one session of the experiment since it was separated from the group during that time.

Immediately after each testing session, the plates as well as all wooden objects were removed from the enclosure. Testing sessions were scheduled at two- to four-day intervals to preserve the subjects’ motivation to interact with the objects.

#### Coding procedure

For coding, the open-source video coding software BORIS (version 9.6.5, release date 02.09.2025; © Olivier Friard) was used. Each session was coded separately for each individual and all manipulations occurring during the 30-min experimental period were coded in detail. For each manipulation, its duration was coded from the subject’s first contact with the object until release (i.e., when the object was dropped). When the start or end of a manipulation could not be directly observed – such as when a subject entered or exited the enclosure with an object – the manipulation duration was recorded from the first or last time the subject was seen. In addition, the manipulated object, the body parts involved, and the behaviors exhibited during object manipulation were coded. These behaviors were classified according to a set of behavioral criteria originally developed for classifying object manipulation in orangutans by Laumer et al. (2025) and subsequently adapted for the present study (see Table B1 in Appendix section B). New behaviors observed in the monkeys were added to the coding scheme as needed.

Moreover, all object-object- and object-substrate-combinations were coded, specifying in each case which object or objects were used.

#### Data analysis

All analyses were done using R (version 4.5.2; R Core Team, 2026) and Rstudio (version 2026.04.0; Posit team, 2026). The *tidyr* (Wickham et al., 2019) and *dyplr* (Wickham et al., 2026) packages were used to structure the raw data and prepare it for statistical analyses (see Appendix section D for detailed information). All figures were created using the ‘ggplot2’ package (Wickham et al., 2019). For statistical analysis, generalised linear mixed models (GLMMs) were fitted. In each model, “Species” (levels: capuchin and patas) was included as a test predictor and “Session” (z-transformed to increase interpretability (Schielzeth, 2010) as control predictor. To avoid issues with pseudoreplication, “Subject” was included as random intercept. For all models, the random slope of session (z-transformed) was included to keep the type I error rate at the nominal level of 0.05 (Barr et al., 2013; Schielzeth & Forstmeier, 2009) (for detailed model structure see SI, section C). Although social rank can be an important source of variation in behavioural studies, it was not determined during data collection and therefore not included as a predictor in the present analyses, as rank is often difficult to assess reliably and consistently and may introduce measurement uncertainty. Age was also not included as a predictor due to the limited sample size. Social rank was assessed by the keepers by means of a questionnaire (see SI, section E) but was not included as a predictor, because rank positions are specific to each group and the sample size did not permit additional predictors. After fitting each model, the potential violations of model assumptions and assessed model stability were evaluated. For each model, it was visually confirmed that the best linear unbiased predictors (BLUPs) per level of the random effects were approximately normally distributed, which was the case for all analyses. The absence of collinearity was validated by calculating the Variance Inflation Factor (VIF) for the models by excluding any interactions and random effects using the package ‘car’ (version 3.0-12; Weisberg & Fox, 2011). For each model, the model stability was assessed by excluding levels of random intercept effects one at a time. Comparing the resulting estimates with those obtained from the model based on all data (Nieuwenhuis et al., 2012), it was determined that all models were of good to moderate stability. For the negative binomial and beta models, the effect of individual fixed effects was tested using the drop1 function, which can be used to compare simpler with more complex models utilizing likelihood ratio tests (Dobson & Barnett, 2002). During the analysis, it was revealed that the subject “Scharti” (capuchin monkey) hardly engaged in object play actions. As she would have been very influential and a clear outlier from the pattern, Scharti was excluded from the “Variability”, “Manipulation Time” and “Number of body parts” analysis. The response variables, error distributions, and deviations from this approach are described below.

**1) Number of smashing behaviour**: Originally, it was intended to fit a GLMM with a Poisson error distribution to inspect differences between species. However, smashing behaviour rarely occurred in patas monkeys, so instead a Fisher’s exact test was applied to test if the number of individuals displaying smashing behaviour differed between species. **2) Variability of object play behaviours**: For the analysis of variability, a generalised linear mixed model (GLMM) with a negative binomial error distribution and logit link function was applied using the function glmer.nb of the lme4 package (Bates et al., 2015). **3) Manipulation time**: To estimate if the total manipulation time differed between species, a GLMM with beta error distribution and logit link function was fitted using the function glmmTMB of the package glmmTMB (Brooks et al., 2017). This approach was selected instead of a linear mixed model because subjects in 44 trials did not show any time manipulating, and including these sessions would have violated the model’s assumptions. By dividing the time spent manipulating by the total observation time, the proportion of time spent manipulating was examined while including data from these trials. Prior to fitting the model, the response was transformed to avoid values being exactly zero or one (Smithson & Verkuilen, 2006). **4) Number of object combinations:** For the analysis of the total number of object combinations (which includes object-object as well as object-substrate combinations), a GLMM with a negative binomial error distribution and logit link function was applied using the function glmer.nb of the lme4 package. Model fitting was performed using the “bobyqa” optimizer with an increased maximum number of iterations to ensure convergence. **5) Number of distinct body parts and tail use during object manipulation:** To analyse if the number of distinct body parts used differed between species, a GLMM with binomial error structure and logit link function was fitted. This approach was used instead of a Poisson model as the number of body parts used has a restricted maximum of seven. By using a binomial model, potential ceiling effects caused by this are naturally accounted for. For this analysis, a two-column response matrix (resp.mat) was used, containing the number of body parts used and the number of body parts not used (Baayen, 2008). Similar to the analysis of the number of smashing behaviour, it was intended to fit a GLMM with a Poisson error distribution to inspect differences between species regarding the number of tail involved in object manipulations. However, the tail was comparatively rarely used in both species, so instead it was analysed whether the ratio of individuals who use their tail and who don’t differs between species by conducting a Fisher’s exact test.

## Results

### 1) Number of smashing behavior

Results of the Fisher’s exact test showed that the number of individuals displaying smashing behavior differed between species (p = 0.01; see Figure 2). Smashing behavior occurred only three times in patas monkeys, twice for Asis and once for Nuru, whereas all capuchin individuals exhibited smashing multiple times throughout the sessions, with the exception of Scharti, who exhibited it only once.

**Figure 2:**
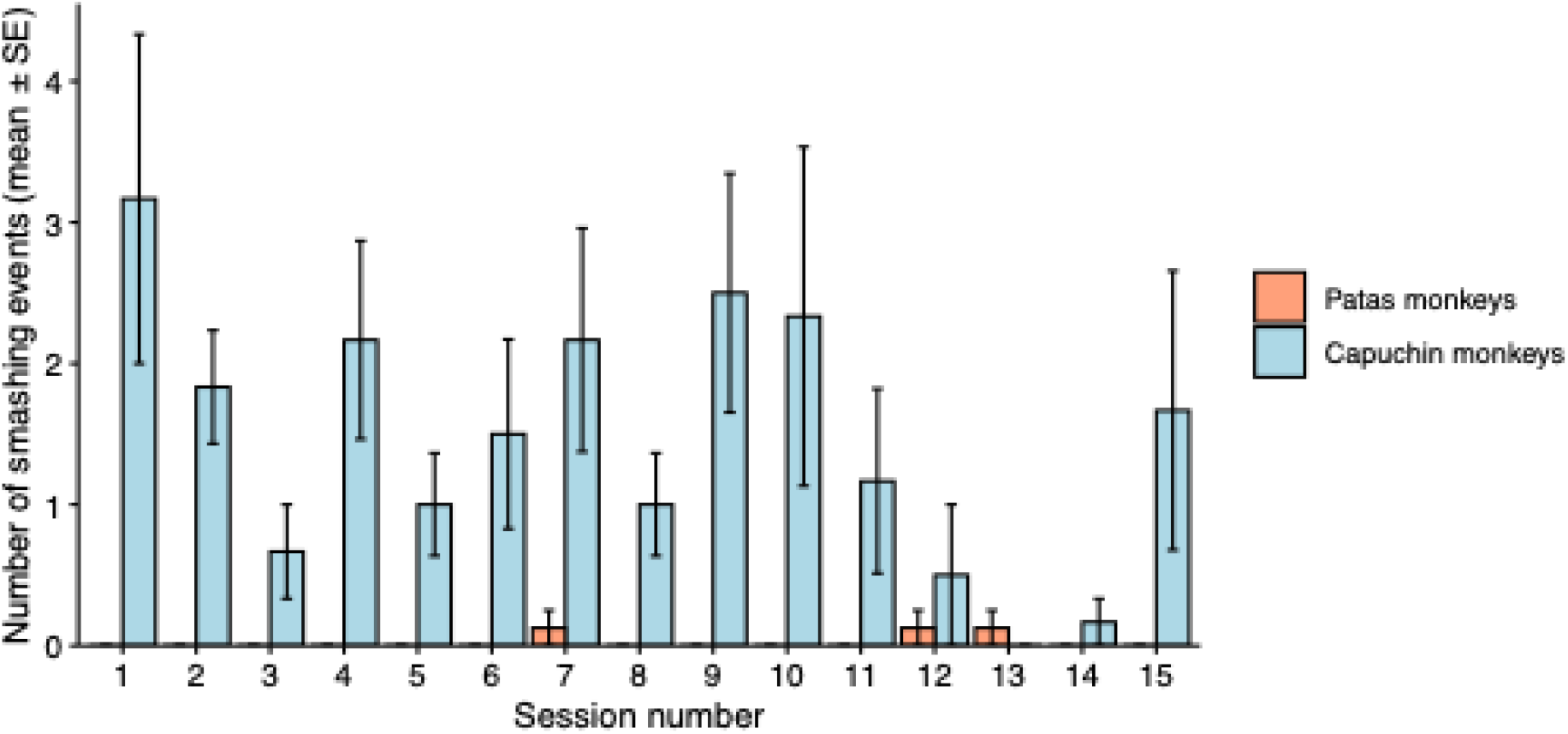
Mean number of smashing events per individual in each session for patas monkeys (n = 8; in orange) and capuchin monkeys (n = 6; in blue). Error bars show the standard error of the mean across individuals.

### 2) Variability of object play behaviors

There was no statistically significant effect of species on variability of exploratory actions (χ2 = 1.033, df = 1, *p* = 0.309; see SI, section D, Figure S3). Both species included individuals showing high behavioral variability, whereas the patas monkeys additionally comprised two individuals characterized by comparatively low counts of variability.

### 3) Manipulation time

There was no effect of species on the proportion of time manipulating (χ2 = 0.01, df = 1, *p* = 0.92; see SI, section D, Figure S4). Four patas monkeys showed comparatively high proportions of manipulation time of 0.3 or higher, whereas the remaining four individuals exhibited much lower proportions. Among the capuchins, one individual showed a comparatively high proportion of manipulation time, while all others had values of approximately 0.2 or lower.

### 4) Number of object combinations

There was no effect of species on the number of object combinations (χ2 = 1.319, df = 1, *p* = 0.251, see SI, section D, Figure S5). Among capuchins, one adult individual showed no object combinations, four individuals (three adults and one juvenile) exhibited low values, and one juvenile individual displayed comparatively higher values. Among the patas monkeys, none of adults performed object combinations. Of the juveniles, two individuals showed a low number of combinations, and two showed a comparatively higher number.

Because not all individuals combined objects, a Fisher’s exact test was additionally applied to test if the number of individuals combining objects differed between species. Similarly to the GLMM, results of the Fisher’s exact test showed that species do not differ in their tendency to combine objects (*p* = 0.11).

### 5) Number of distinct body parts and tail use

There was no effect of species on the number of body parts used (χ2 = 0.289, df = 1, *p* = 0.59, see SI, section D, Figure S6). Fitted values indicated that capuchin monkeys consistently used at least three body parts during object manipulation. In patas monkeys, three individuals showed lower fitted values, whereas the remaining five individuals used four or more body parts.

The results of the Fisher’s exact test showed that there was no effect on the ratio of individuals who use their tail and who don’t between the species (*p* = 0. 277, see SI, section D, Figure S7). The tail was rarely used by patas monkeys, occurring only once in two sessions of Atu and once in a session of Nuru. In contrast, four capuchin individuals used their tail at least once during object manipulation, with only Tica using it several times across multiple session.

## Discussion

Both species manipulated the objects and both displayed a range of behaviors, some of which can be classified as secondary combinatory manipulations that combine objects with other objects or with substrates. Although species-specific differences related to ecological variation were expected, statistically significant differences were only found in the number of subjects exhibiting smashing behavior during object manipulation, with smashing occurring frequently in all capuchins except one individual, but very rarely in patas monkeys. The predictions concerning behavioral variability, object combinations, manipulation time, and tail use were not supported.

### 1) Smashing behavior

All six capuchins smashed objects, five of them repeatedly, whereas only two of eight patas monkeys did so, on three occasions in total. This difference is consistent with the large repertoire of extractive foraging techniques documented in wild capuchin populations, including the repeated cracking of nuts, seeds, fruits, and cacti as well as the digging for high-carbohydrate underground tubers and protein-rich insects (e.g., de A. Moura & Lee, 2004; Falótico & Ottoni, 2016; Fragaszy et al., 2005; Fragaszy & Boinski, 1995; Izawa & Mizuno, 1977; Mannu & Ottoni, 2009; Melin et al., 2014; Moura, 2007; Ottoni & Mannu, 2001; Perry & Ordonez Jiménez, 2006; Rose, 1994; Souto et al., 2011; Struhsaker & Leland, 1977). Patas monkeys feed mainly on items that require little mechanical processing, such as parts of *Acacia* plants, fruits of other savanna trees and small arthropods such as ants (Isbell, 2013; Kingdon, 1997) The finding parallels results obtained with the same protocol in birds, in which the type of object manipulation reflected foraging ecology: insertions predominated in New Caledonian crows, which extract food with tools, and play caching in ravens, which cache food (Auersperg et al., 2014). Smashing is not specific to species that use tools, however. Captive kea, which are not known to use tools habitually, also smash objects on substrates (Gajdon et al. (2014), as cited in Auersperg et al. (2015)). Smashing during play is therefore best interpreted as an expression of foraging ecology, and the present data do not show that it contributes to the development of tool use.

### 2) Behavioral variability

Contrary to our prediction, that capuchins would show higher behavioral variability due to higher ecological flexibility (Fedigan, 2017) than Patas monkeys that occupy a narrower ecological niche, behavioral variability did not significantly differ between species. Both species included highly variable individuals, and the patas group additionally included two individuals with low variability. In parrots, the yellow-billed Amazon (*Amazona collaria*), which occupies a range of habitats combined objects at low rates, whereas the closely related black-billed Amazon (*A. agilis*), which is restricted to one habitat manipulated the objects frequently but never combined them (Auersperg et al., 2015). Ecological breadth may thus be expressed in the complexity of object combinations rather than in the diversity of object actions. Age composition may also have influenced the comparison. In the patas group, juveniles showed the highest variability. Perry (2020) reported that younger white-faced capuchins (*Cebus capucinus*) showed a more diverse behavioral repertoire than adults in extractive foraging tasks, so the proportion of juveniles in a group may effect variability at the group level. The capuchin group in present study included two juveniles alongside four adults, while the patas group included four juveniles alongside four adults.

### 3) Manipulation time

Contrary to our prediction, capuchin monkeys did not spend significantly more time manipulating objects than the patas monkeys. Cenni et al. (2024) argued that differences relevant to the acquisition of tool use may lie in qualitative aspects of manipulation rather than total time spent in object play. This pattern is markedly reflected in the present study, where significantly more capuchins than patas monkeys exhibited smashing behavior. In addition, the objects used in this study offered no food reward. Extractive foraging demands persistence (Branca et al., 2025; Ibáñez de Aldecoa et al., 2024), and such persistence may be expressed when individuals are faced with ecologically relevant challenges that involve immediate rewards, such as food extraction. In patas monkeys, juveniles showed the highest proportion of manipulation time, consistent with the decline of play with age (Pellis & Ham, 2024). In capuchins, one juvenile showed the highest and the other the lowest proportion, which may be explained by individual variation within the species rather than a systematic age-related pattern.

### 4) Object combinations

The number of object combinations did not significantly differ between species. Five out of six capuchins and four out of eight patas monkeys combined objects, and the highest individual values occurred in both species. Combination rates were low overall. The estimated mean for capuchins was 0.43 and for patas 0.12 combinations per individual and session. Under the same protocol, young Goffin cockatoos combined two objects 4.6 times per session on average, while young New Caledonian crows combined objects more frequently than ravens and jackdaws (Auersperg et al., 2014, 2015). The avian pattern was therefore not replicated here. This may reflect differences between taxa in combinatory play, differences in motivation, or features of the setup. However, the observed discrepancy may indicate species-specific differences in the expression of combinatory object play, differences in motivation, or features of the setup. The low number of object combinations in capuchins may be explained by the spatial arrangement of the activity plates. Given the predominantly arboreal lifestyle of capuchins (Havmøller et al., 2021), the activity plates located on the floor possibly did not represent an attractive location for prolonged engagement. Although not formally quantified, it was repeatedly observed during testing that capuchins collected objects from the enclosure floor and immediately carried them to elevated climbing structures, thereby moving them away from the activity plates. This likely reduced opportunities for object-substrate interactions, resulting in a generally low number of total object combinations. Future research could address this challenge by placing the experimental apparatus in elevated locations to better align with species-typical spatial preferences.

Object combinations occurred in juvenile patas monkeys, whereas three of four adult capuchins combined objects. Age was not included in the models because of the small sample, so these patterns are descriptive. Consistent with the prediction that juveniles of both species performing more object combinations than adults, object combinations in patas monkeys exclusively occurred in juveniles. Interestingly, Njabi, a four-year-old patas male currently transitioning into adulthood, exhibited the lowest number of object combinations among all juveniles. While this may in part reflect individual variation, it could also be the consequence of a developmental decline in general play and also object play behavior as puberty approaches, a trend also reported in humans and some birds (e.g., Bapat et al., 2025; Miller et al., 2015; Pellegrini, 2009). Evidence from birds shows that adult object play is not a general feature of species that use tools. Adult New Caledonian crows showed almost no interest in the objects, whereas adult Goffin cockatoos combined objects as often as juveniles (Auersperg et al., 2014). The authors related persistent adult play in Goffin cockatoos to their generalist diet and use of diverse habitats. Object combinations in adult capuchins would be consistent with this interpretation, because capuchins are also dietary generalists (Fedigan, 2017). With the four adult capuchins, however, this remains a descriptive observation.

### 5) Body parts and tail use

As expected, the species did not differ in the number of body parts used during object manipulation events. This is consistent with the findings by Parker (1978), who found broadly similar responses to a nylon rope in capuchins and guenons, which belong to the same subfamily as patas monkeys, whereas great apes used more body parts.

Contrary to our prediction, the proportion of individuals that used their tail during object manipulation did not differ between species. This is surprising since capuchins rely more on tail use due to their prehensile tail, which is adapted for grasping and can act as a fifth extremity. However, the evolutionary advantage of the prehensile tail is likely not primarily its role in complex object manipulation, but rather its function as an adaptation to an arboreal lifestyle. Evidence for this was provided by Naas et al. (2025) who found that *Sapajus apella* individuals in the rainforest on the border between Brazil and French Guiana predominantly employ their prehensile tail to anchor themselves and maintain stability in above-branch positions. While the tail thus primarily serves a postural function, the main body parts involved in object manipulation in primates remain the hands (Crast et al., 2009; Marzke & Wullstein, 1996; Parker, 1974).

#### Limitations

The findings are limited by a small sample size (capuchins: n = 6 and patas: n = 8) and a high degree of individual variation. Each category was represented by a single species, so the differences may reflect phylogeny, ecology, or group history rather than tool use. The groups also differed in age composition and in previous experimental experience. Neither age nor social rank could be included in the analysis. Following the original protocol, the subjects were tested in their social group, where group dynamics and social rank may have influenced the present results. This is particularly the case in the capuchin group, where the lowest-ranking individual Scharti rarely engaged in object play, possibly due to avoidance of object interaction in the presence of higher-ranking individuals. The coatis that shared the capuchin enclosure were near the activity plates in seven of the 15 sessions. Although they did not interact with the objects beyond sniffing, an effect of their presence on the capuchins’ use of the plates cannot be excluded. Although considerable effort was made to record all object interactions, continuous observation of all subjects was not always feasible, when multiple individuals were manipulating objects simultaneously in different areas of the enclosure, which happened rarely. As a result, it is possible that not all object manipulations were captured. Finally, captivity may elevate exploratory manipulation. In a recent study by Laumer et al. (2025) zoo-housed Sumatran orangutans showed significantly higher rates of exploratory object manipulation compared to wild conspecifics, possibly due to the absence of predators and reduced foraging demands, which allows more time and energy for exploring the environment and manipulating objects (Haslam, 2013; Kummer & Goodall, 1985). Furthermore, Laumer et al. (2025) observed that captive orangutans used more body parts during exploration, exhibited a larger repertoire of exploratory actions overall and have a higher tendency to explore multiple objects simultaneously, including tool use, as enrichment in zoo settings may enable more intricate and sustained combinatory manipulations than in natural environments (Milgram et al., 2006; Puteri & Ruppert, 2017; Scheel, 2018; Tarou et al., 2004; Tripp, 1985). Although orangutans were not part of the present study, these findings are nevertheless relevant and may also help explain why differences between patas monkeys and capuchins in manipulation time and number of object combinations were less pronounced than initially expected, and why variability in object play behavior in patas monkeys tended to be higher than predicted.

## Conclusion

Capuchins and patas monkeys differed in one specific form of manipulation, smashing, but not in behavioral variability, manipulation time, number of body parts used, tail use, or the number of object combinations. As in birds tested with the same protocol, the form of manipulation reflected foraging ecology. The prediction that a species that uses tools shows more complex combinatory play was not supported. To our knowledge this is the first application of this protocol in primates. Testing whether combinatory object play is associated with tool use in primates will require larger samples, balanced age compositions and several species per category.

## Author contributions

IBL and AMIA conceptualized the study. TB collected the data. RF analyzed the data. TB wrote the first draft and IBL and AMIA edited the paper.

## Acknowledgements

We would like to thank Rostock Zoo, especially the curator of mammals Daniela Lahn and the two zoo keepers Katrin Reimer and Ines-Chatrin Tischler for permission and support in conducting this research. Additionally, we thank Prof. Dr. Stefan Richter (Department of General and Systematic Zoology, University of Rostock) for his advice and feedback.

## Data Availability

Data and analysis scripts will be uploaded during the submission process and upon publication the data will be shared publicly as supplementary materials.

