## Supplementary Material for "Percussive manipulations distinguish object play in capuchin and patas monkeys"

### (A) Additional information on the experimental procedure

#### Habituation phase

Activity plates were introduced into the indoor enclosures of patas and capuchin monkeys several days before the experiments. The aim was for each individual to interact with or approach the plates at least once, thereby minimizing the likelihood of neophobic behavior during the experimental sessions later on.

However, the plates were removed from the capuchin enclosure after 24 hours and from the patas enclosure after 72 hours (first habituation phase), as the animals had started to damage the apparatus. Following this, the plates were repaired and reinforced to enhance their stability. They were then reintroduced into the enclosures under supervision, with access limited to approximately 90 minutes over the course of four days during the second habituation phase.

During both habituation phases, the patas monkeys showed very little evidence of neophobia: all eight individuals approached and touched the activity plates within 24 hours during the first habituation phase and again by the second session of the second habituation phase. In contrast, the capuchin monkeys required more time, with all individuals having interacted with the activity plates only by the third session of the second habituation phase.

### Apparatus and Experimental setup

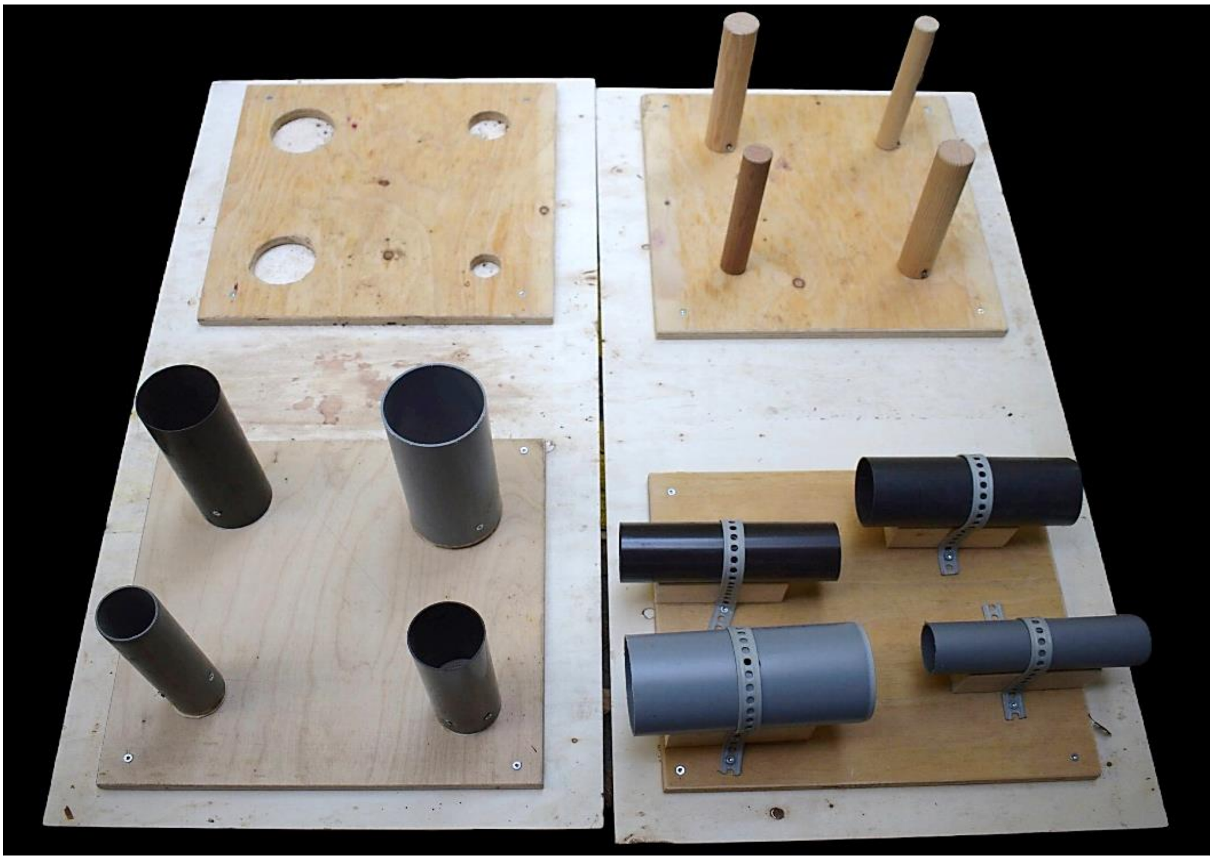

**Figure S1**: Image of the activity plates according to the dimensions specified in the methods section.

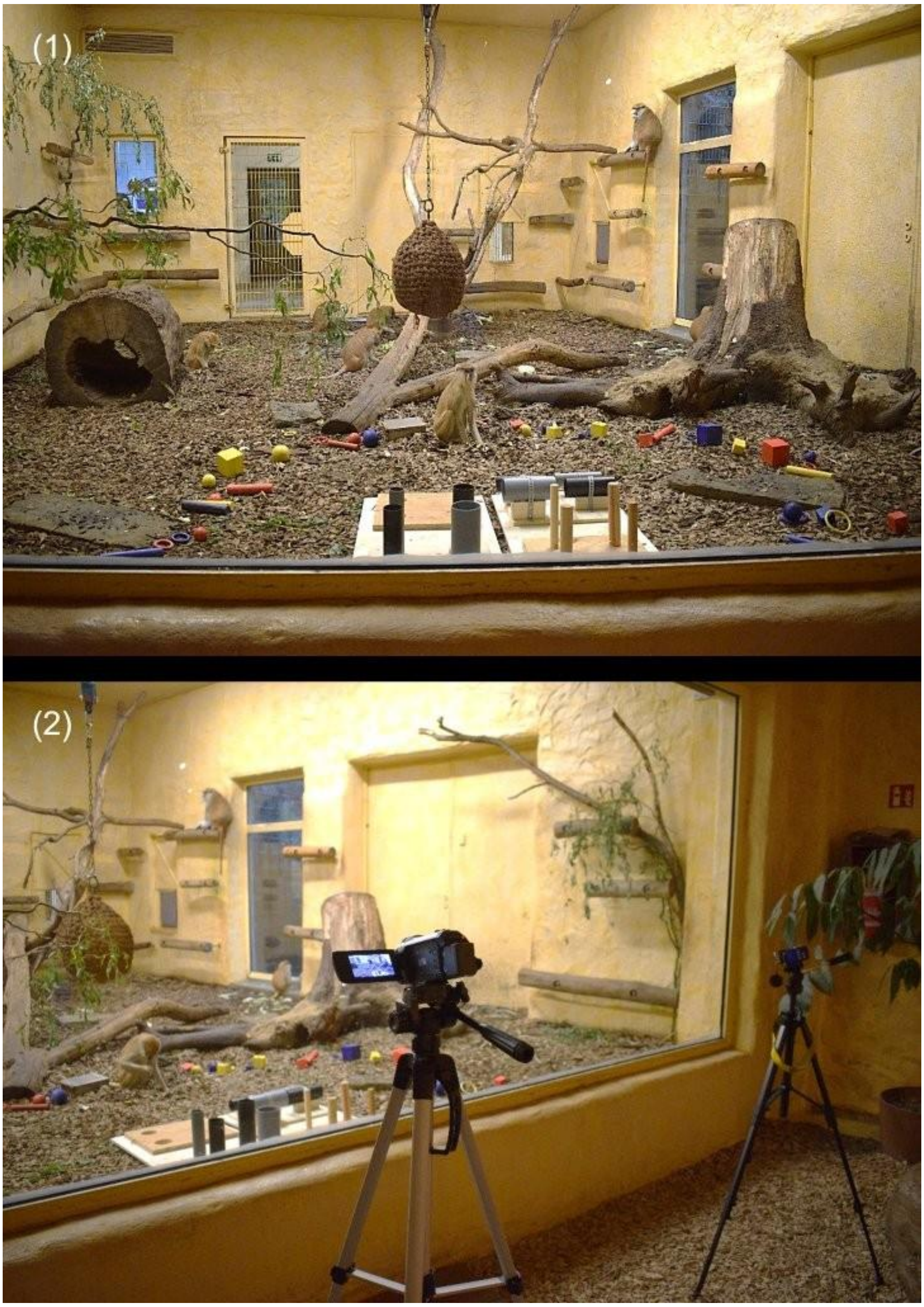


**Figure S2**: Representative images of the experimental setup used during testing sessions with the patas monkeys. An identical configuration was employed for the testing sessions involving the capuchin monkeys. **(1)** Position of activity plates and wooden objects in the enclosure. **(2)** Position of video cameras.

### Testing schedule and constraints

The experiments were conducted early in the mornings before the official opening hours of the zoo to avoid distraction or interruption by visitors. During the experiments, the animals were intended to remain in the indoor enclosure containing the apparatus. However, due to weather conditions and internal logistical constraints, including occasional limitations in relocating animals to the indoor enclosure prior to the experiments, it was not always feasible to maintain this restriction. As a result, animals occasionally had access to both the indoor and outdoor enclosure area, allowing free movement between these spaces during some experimental sessions.

Due to the presence of white-nosed coatis in the capuchin monkeys' enclosure, the experimental procedure for the capuchins was slightly adjusted. Whenever the coatis were in the vicinity (approximately 50 cm radius) of the activity plates, thereby potentially hindering the capuchin monkeys to approach, the duration of their proximity was documented. These time intervals were then added to the 30-minute observation period to ensure that each trial comprised a total of 30 minutes of observation without interference. This procedure was implemented to minimize potential confounding effects, as the focal individuals might have been discouraged from approaching the activity plates while the coatis were nearby.

### (B) Additional information on the coding process

**Table S1**: List and descriptions of all behaviors exhibited during object manipulation and used for the coding procedure.

| **Behavior** | **Description** |
|---|---|
| Balancing on body part | Adjust or reposition an object on the body |
| Biting | Cutting an object in the mouth with teeth |
| Biting off | Using the teeth to bite off a piece of the object |
| Bouncing | Lifting an object (especially a ball) a short distance and releasing it so it drops; lift-and-drop action is repeated multiple times |
| Carrying around | Transporting an object (directed movement) |
| Chewing | Repeatedly biting on an object that is inside the mouth |
| Collecting | Collecting several objects |
| Construct | Bring together two or more objects or an object with a substrate |
| Dragging | Pulling an object over substrate (usually the ground) without letting go of it |
| Dropping | Make an object fall by letting go of it intentionally |
| Gnawing | Repeatedly biting on an object that is not inside the mouth |
| Grid Insertion | Partially or fully inserting an object through the grid/ bars/ wire mesh |
| Inserting | Placing an object into a tube or a hole on the activity plates |
| Knocking | Gently (and repeatedly) hitting with knuckles at a surface/ object |
| Knocking over | Strike or collide an object so as to cause them to fall to the ground |
| Laying / standing / sitting in | Laying, standing or sitting in or on the object |
| Licking | Passing tongue over object |
| Lifting | Lifting up an object and then replacing it where it came from |
| Looking through | Looking fixedly through an object |
| Pulling | Dragging an object that is being held by another subject or is stuck somewhere |
| Pushing on surface | Putting an object on a surface (i.e., a wall) and holding it with one or two hands, thereby not moving the object any further |
| Putting on head | Placing an object onto own head |
| Reaching into | Sticking body part (hand, arm, foot, head) into a substrate or object (e.g., tree hole or big piece of dead wood) |
| Retrieving object | Taking an object out of a closed or enclosed structure (i.e., a tube) using the hands or feet |
| Ringstacking | Putting ring-shaped objects onto a wooden pole on the activity plates or onto any tapered end of a climbing structure |

| | |
|---|---|
| | (i.e., the end of a branch) |
| Rolling object | Moving or turning the object on a surface without lifting |
| Rolling on body part | Letting an object roll from the hands or feet towards the body along the arms or legs |
| Rotating | Moving or rotate an object in a circular motion (distinct from rolling) |
| Rubbing object against body part | Moving object against a body part repeatedly (slow or fast) while applying pressure (different to sliding which is a singular motion) |
| Rubbing on surface | Moving an object on the surface repeatedly (slow or fast) while applying pressure (different to sliding which is a singular motion) |
| Slapping | Gently hitting an object with the palm of the hand |
| Sliding over surface | Pulling an object over another object or substrate in a sliding motion |
| Smashing | Hitting an object with force against a surface/ that is laying on a surface |
| Smashing object with object | Hitting an object with another object. Often repeatedly. |
| Smelling | Sniffing an object |
| Tearing apart | Pulling objects into two pieces by applying force |
| Throwing | Propel object with force through the air |
| Tipping | Touching an object repeatedly with the fingertips, without moving it any further |
| Touching | Deliberately touching an object with hands or feet, thereby looking at it intently, but not performing any other actions |
| Transferring | Handing an object from one body part to another |
| Turning around | Turning object around in hands and/ or feet. Often includes repeated handing of objects from hands to feet and back → Transferring |

### (C) Additional information on the statistical analysis

**Table S2:** List and description of variables calculated from the raw data after coding. Additionally, the corresponding excel file included additional information on the subjects, such as age, species, sex, and the experimental session. This processed dataset was subsequently used for all statistical analyses.

| **Variable** | **Description** |
|---|---|
| Variability | The number of unique object play behaviors a subject displayed within one session.<br><br>Example: If subject A exhibited the behavior "touching" three times, "smashing" twice and "dropping" once within one session, the variability for subject A would be 3 (as there were three unique behaviors, regardless of frequency). |
| Total manipulation time | The total time (in seconds) per session that a subject spent manipulating objects. |
| Number of smashing | The total count of smashing behaviors exhibited by a subject within one session. |
| Number of body parts | The number of different body parts a subject used during object play within one session.<br><br>Example: If subject A used the left hand twice, the right hand twice, and the feet once, the number of body parts would be 3 (as three distinct body parts were used). |
| Number of tail use | The total count of tail use during object play per subject and session. |
| Number of object-object-combinations | The total count of object-object combinations performed by a subject within one session. |
| Number of object-substrate combinations | The total count of object-substrate combinations performed by a subject within one session. |
| Number of total combinations | The total count of object combinations (sum of the number of object-object and object-substrate combinations) performed by a subject within one session. |

### Model Structure

Variability:

```
lme4::glmer.nb(Variability ~ Species + z.Session + (1 + z.Session | Subject), data = data)
```

Manipulation time:

```
glmmTMB(tr.Perc.ManipulationTime ~ Species + z.Session + (1 + z.Session || Subject), data = data, family=beta_family)
```

Number of object combinations:

```
lme4::glmer.nb(Number_of_comb_total ~ Species + z.Session + (1 | Subject), data=data, control = glmerControl(optimizer =
"bobyqa", optCtrl = list(maxfun = 200000)))
```

Number of distinct body parts used during object manipulation:

```
glmer(resp.mat ~ Species + z.Session + (1 + z.Session | Subject), data = data, family = binomial)
```

### (D) Additional results

#### Variability of object play behaviours

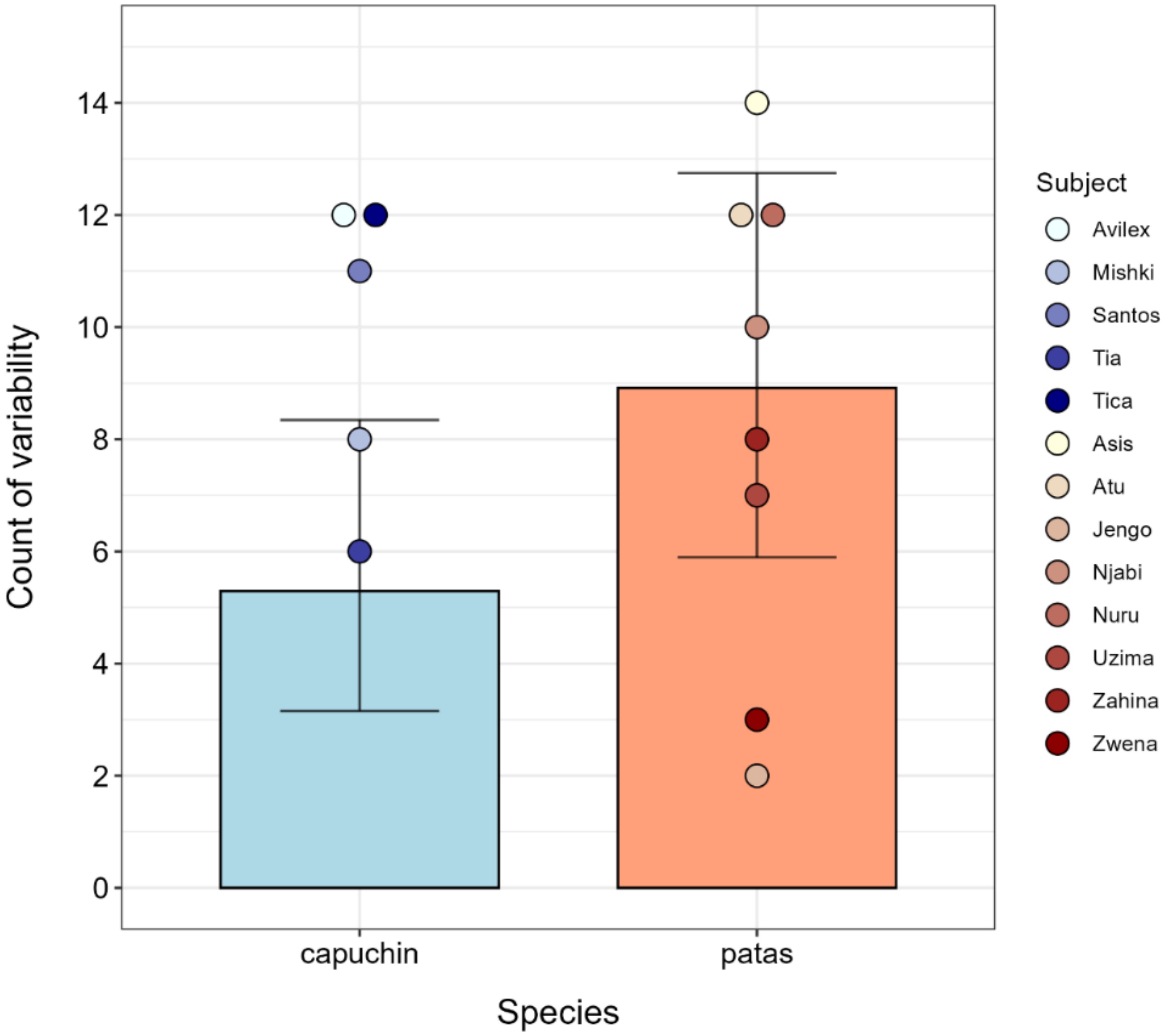


**Figure S3:** Model-based estimated counts of object play behavior variability per species. Bars represent model-fitted values. Error bars indicate 95% confidence intervals. Colored points indicate the median variability per subject across sessions (capuchins: N = 5 and patas: N = 8).

### Manipulation time

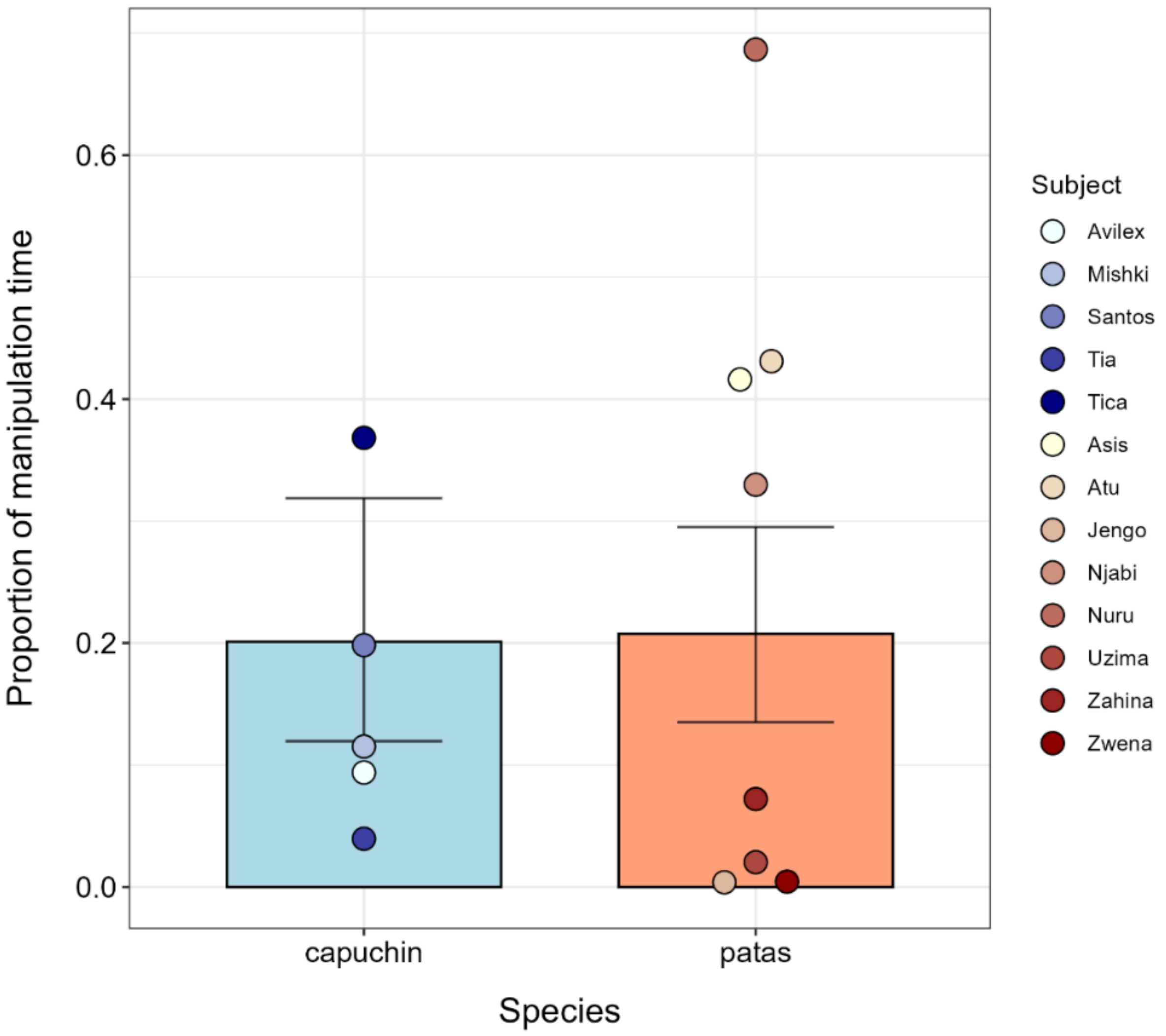


**Figure S4:** Model-based estimates of the proportion of time spent manipulating objects per species. Bars represent fitted values. Error bars indicate 95% confidence intervals. Colored points represent the subject-level median proportion across sessions (capuchins: N = 5 and patas: N = 8).

### Number of object combinations

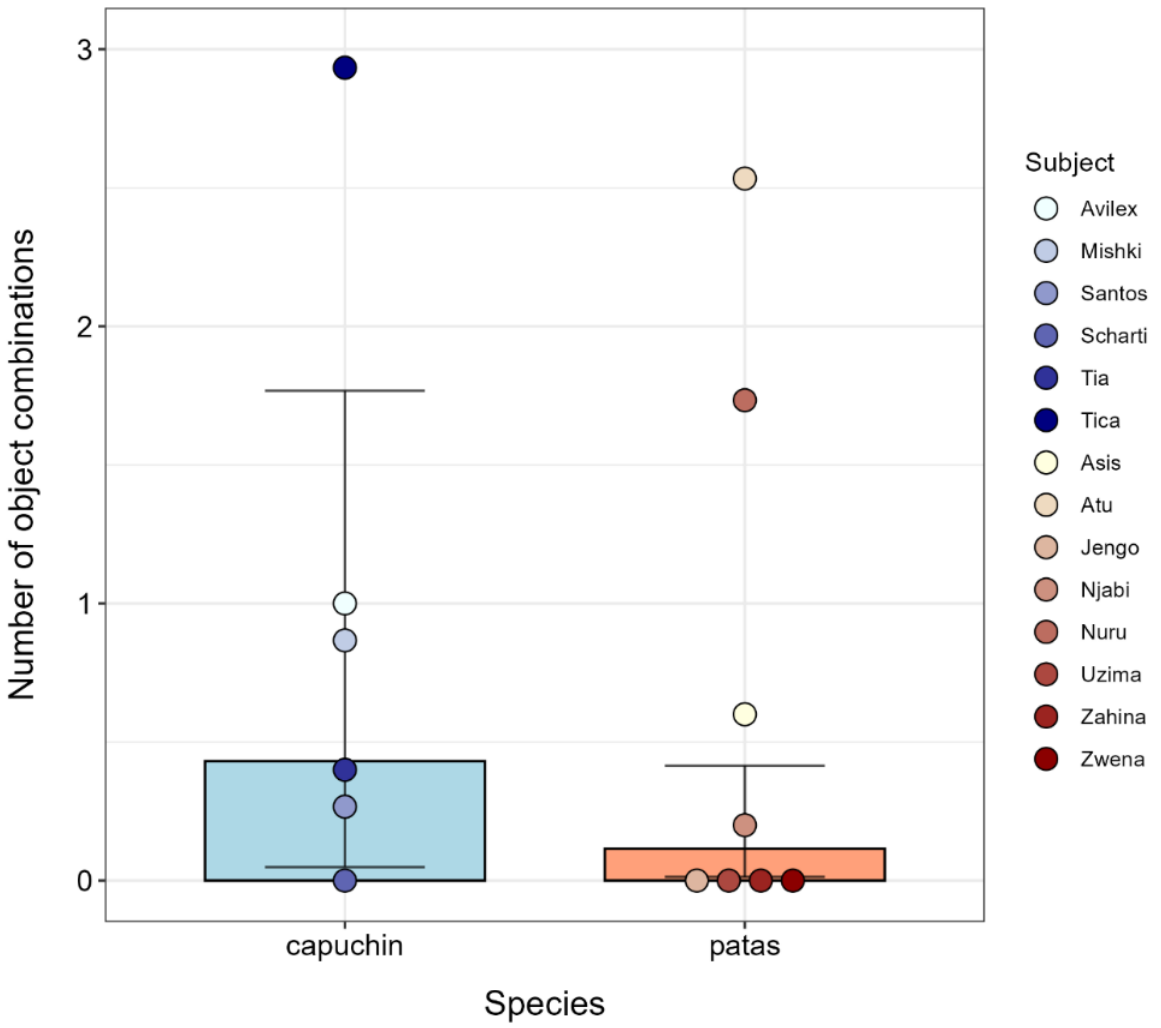


**Figure S5:** Model-based estimates of the number of object combinations (sum of object-object and object-substrate combinations) per species. Bars represent fitted values. Error bars indicate 95% confidence intervals. Colored points indicate the median number of object combinations per subject across sessions (capuchins: N = 6 and patas: N = 8).

### Number of distinct body parts used during object manipulation

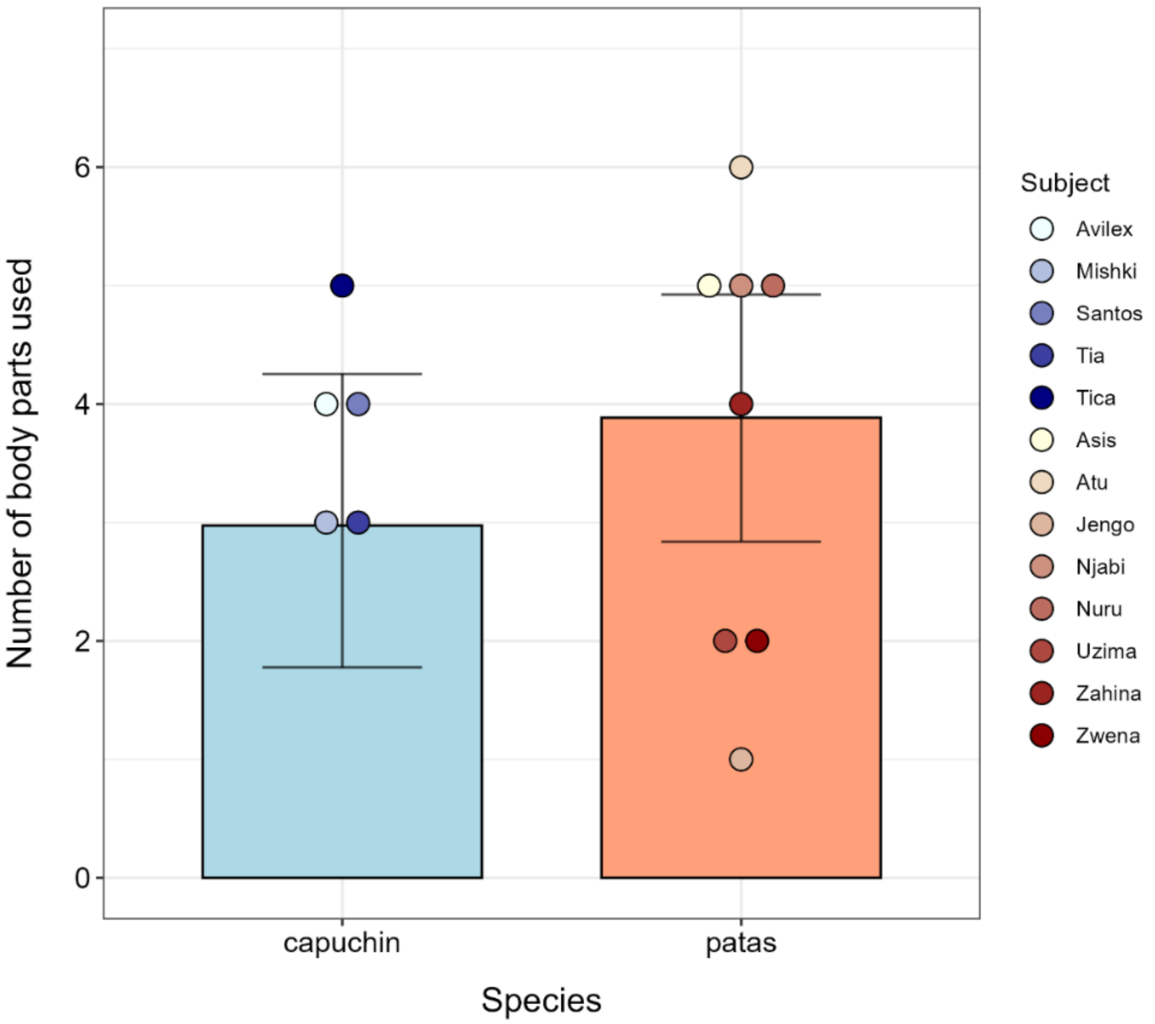


**Figure S6:** Model-based estimates of the number of body parts used during object manipulation events per species. Bars represent fitted values. Error bars indicate 95% confidence intervals. Colored points indicate the median number of body parts used during object manipulation per subject across sessions (capuchins: N = 5 and patas: N = 8).

### Number of tail use during object manipulations

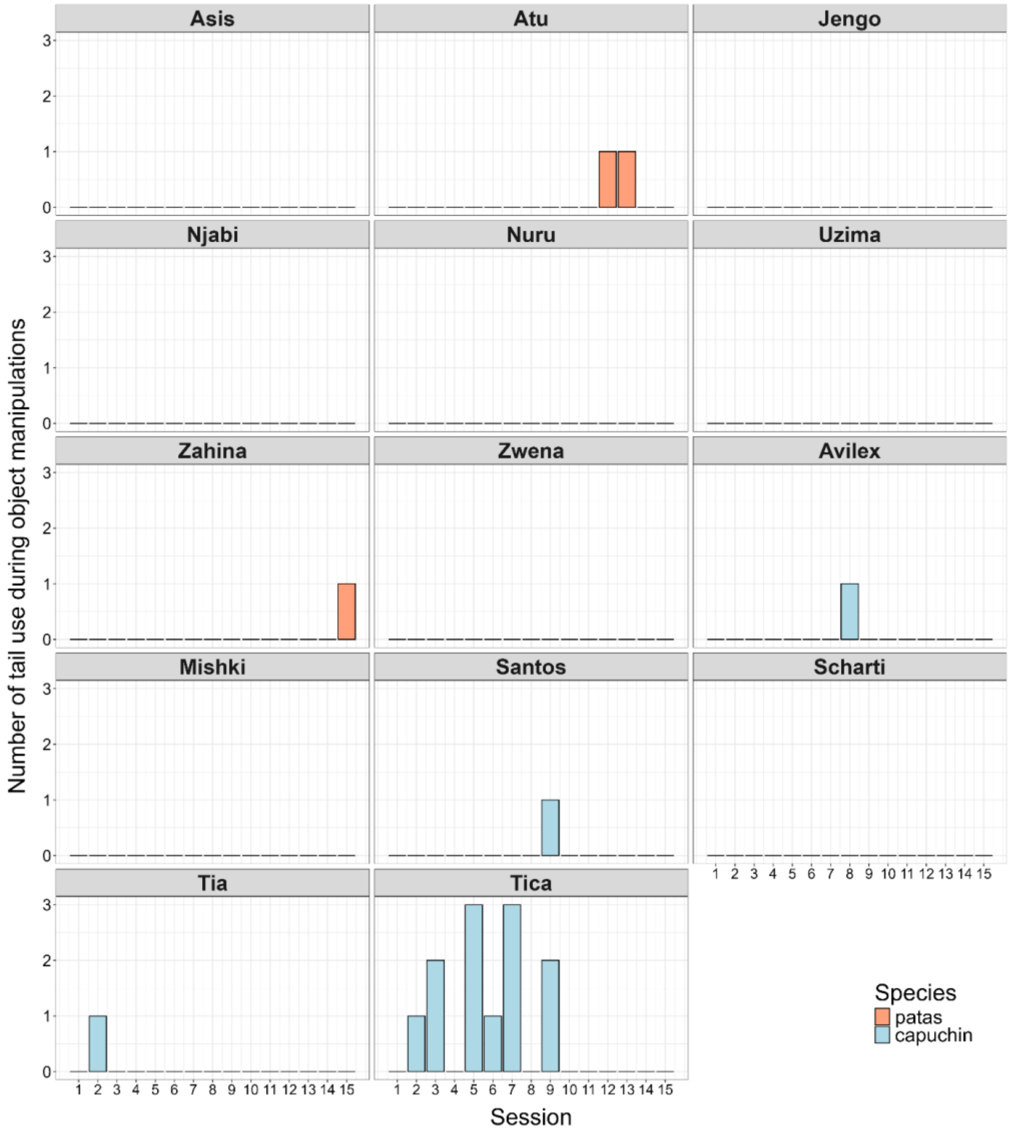


**Figure S7:** Number of times the tail was used during object manipulation per session for each subject.

### (E) Animal assessment questionnaires

**<u>Questionnaire 1</u>**

**Keeper's name: C. Tischler** **Date: 04.01.2026**

- **Dominance hierarchy**

| **Patas monkeys** | **Capuchin monkeys** |
|---|---|
| **1.** Zahina | **1.** Santos |
| **2.** Zwena | **2.** Mishki |
| **3.** Njabi | **3.** Tica |
| **4.** Atu | **4.** Tia |
| **5.** Nuru | **5.** Avilex |
| **6.** Asis | **6.** Scharti |
| **7.** Jengo | |
| **8.** Uzima | |

- **Have you ever observed tool use in the patas monkeys (beside my experiments)? If so, which individuals used tools and what did they do exactly?**

No, I have never observed tool using behaviour in our patas monkeys.

- **Have you ever observed tool use in the capuchin monkeys (beside my experiments)? If so, which individuals used tools and what did they do exactly?**

Santos and Avilex sometimes collect and prepare sticks laying around the enclosure and use them to fish for food or to poke around in crevices and cracks. Tica has been observed while poking with sticks as well.

**How curious is the animal on a scale of 1 to 10?**
**(1= not curious at all | 10 = very curious)**

| **Patas monkeys** | **Capuchin monkeys** |
|---|---|
| **Asis:** 7 | **Avilex:** 8 |
| **Atu:** 7 | **Mishki:** 7 |
| **Jengo:** 8 | **Santos:** 8 |
| **Njabi:** 8 | **Scharti:** 6 |
| **Nuru:** 8 | **Tia:** 8 |
| **Uzima:** 2 | **Tica:** 8 |
| **Zahina:** 5 | |
| **Zwena:** 5 | |

**How anxious/nervous is the animal on a scale of 1 to 10?**
**(1= not anxious/nervous at all | 10 = very anxious/nervous)**

| Patas monkeys | Capuchin monkeys |
|---|---|
| **Asis:** 3 | **Avilex:** 5 |
| **Atu:** 3 | **Mishki:** 5 |
| **Jengo:** 9 | **Santos:** 5 |
| **Njabi:** 5 | **Scharti:** 5 |
| **Nuru:** 5 | **Tia:** 5 |
| **Uzima:** 10 | **Tica:** 5 |
| **Zahina:** 7 | |
| **Zwena:** 7 | |

For both species, anxiousness depends very much on the situation. Examples for situations where all animals show a high degree of anxiousness in general are: veterinary treatments, high noise exposure due to jet aircraft overflight and new enclosure equipment.

**For how long have you known the animals?**

| Patas monkeys | Capuchin monkeys |
|---|---|
| **Asis:** since birth (1 year) | **Avilex:** since birth (22 years) |
| **Atu:** since birth (2 years) | **Mishki:** 15 years |
| **Jengo:** 9 years | **Santos:** 16 years |
| **Njabi:** since birth (4 years) | **Scharti:** since birth (34 years) |
| **Nuru:** since birth (3 years) | **Tia:** since birth (3 years) |
| **Uzima:** since birth (19 years) | **Tica:** since birth (5 years) |
| **Zahina:** since birth (9 years) | |
| **Zwena:** since birth (10 years) | |

**<u>Questionnaire 2</u>**

**Keeper's name: K. Reimer** **Date: 13.12.2025**

**Dominance hierarchy**

| Patas monkeys | Capuchin monkeys |
|---|---|
| **1.** Zahina | **1.** Santos |
| **2.** Zwena | **2.** Mishki |
| **3.** Njabi | **3.** Avilex |
| **4.** Nuru – ranking changes constantly (3.–5.) | **4.** Tica – ranking changes constantly (3.–5.) |
| **5.** Atu | **5.** Tia |
| **6.** Asis | **6.** Scharti |
| **7.** Uzima | |
| **8.** Jengo | |

**Have you ever observed tool use in the patas monkeys (beside my experiments)? If so, which individuals used tools and what did they do exactly?**

All patas monkeys except Jengo sometimes rub bark or sticks against other branches or the walls. Juveniles (Atu, Asis and Nuru) imitate this behaviour and occasionally rub sticks against their chests.
[Note: according to the definition used in the present study, these behaviours do not represent tool use]

**Have you ever observed tool use in the capuchin monkeys (beside my experiments)? If so, which individuals used tools and what did they do exactly?**

All of the capuchin monkeys use stones to crack open nuts, thereby smashing the nut on the stone. Santos, Tica, Tia, and Avilex crack open coconuts by smashing them against the wall or enclosure structures. Furthermore, they use small sticks to poke around in holes and cracks.

**How curious is the animal on a scale of 1 to 10?**
**(1= not curious at all | 10 = very curious)**

| Patas monkeys | Capuchin monkeys |
|---|---|
| **Asis:** 9 | **Avilex:** 10 |
| **Atu:** 9 | **Mishki:** 10 |
| **Jengo:** 6 | **Santos:** 10 |
| **Njabi:** 10 | **Scharti:** 10 |
| **Nuru:** 10 | **Tia:** 10 |
| **Uzima:** 6 | **Tica:** 10 |
| **Zahina:** 9 | |
| **Zwena:** 9 | |

**How anxious/nervous is the animal on a scale of 1 to 10?**
**(1= not anxious/nervous at all | 10 = very anxious/nervous)**

All patas monkeys display minimal anxiety (score of 1) towards familiar objects and enclosure equipment, but extreme anxiety (score of 10) towards unfamiliar ones.

It is difficult to quantify behaviour using a numerical scale in capuchin monkeys. They exhibit anxiety by vocalising and fleeing in response to noise that they are unable to identify, but in general, they are not very anxious.

**For how long have you known the animals?**

| Patas monkeys | Capuchin monkeys |
|---|---|
| **Asis:** since birth (1 year) | **Avilex:** since birth (22 years) |
| **Atu:** since birth (2 years) | **Mishki:** 15 years |
| **Jengo:** 9 years | **Santos:** 16 years |
| **Njabi:** since birth (4 years) | **Scharti:** since birth (34 years) |
| **Nuru:** since birth (3 years) | **Tia:** since birth (3 years) |
| **Uzima:** since birth (19 years) | **Tica:** since birth (5 years) |
| **Zahina:** since birth (9 years) | |
| **Zwena:** since birth (10 years) | |
